# Neuronal NADPH oxidase 2 in Parkinson’s disease pathogenesis

**DOI:** 10.1101/2021.05.13.443999

**Authors:** Matthew T. Keeney, Eric K. Hoffman, Kyle Farmer, Christopher R. Bodle, Marco Fazzari, Alevtina Zharikov, Sandra L. Castro, Julia K. Kofler, Eugenia Cifuentes-Pagano, Patrick J. Pagano, Edward A. Burton, Teresa G. Hastings, J. Timothy Greenamyre, Roberto Di Maio

## Abstract

Mitochondrial dysfunction and oxidative stress are strongly implicated in the pathogenesis of Parkinson’s disease (PD) and there is evidence that mitochondrially-generated superoxide can activate NADPH oxidase 2 (NOX2), which is a major enzymatic generator of superoxide. Although NOX2 has been examined in the context of PD, previous studies have focused on microglial function; the role of neuronal NOX2 in PD pathogenesis remains to be defined. Here we devised and validated a proximity ligation assay for NOX2 activity and demonstrated that in human PD and 2 models thereof, neuronal NOX2 is highly active in substantia nigra dopamine neurons. Further, NOX2 activity is responsible for accumulation, post-translational modification and oligomerization of α-synuclein as well as activation of leucine-rich repeat kinase 2 (LRRK2). Administration of a brain-penetrant, specific NOX2 inhibitor prevented NOX2 activation and its downstream effects *in vivo* in a rat model of PD. We conclude that neuronal NOX2 is a major contributor to oxidative stress in PD, to α-synuclein pathology and to LRRK2 activation in idiopathic PD. In this context, NOX2 inhibitors hold potential as a disease-modifying therapy in PD.

**Summary:** In dopamine neuron, NADPH oxidase isoform 2 amplifies the oxidative stress-related pathogenic cascade in Parkinson’s disease

## Introduction

The pathogenesis of Parkinson’s disease (PD) is multifactorial and the exact mechanisms accounting for selective nigrostriatal cell loss are still incompletely understood. Oxidative stress has long been implicated in PD (1) and there is consensus on the exquisite vulnerability of dopamine (DA) neurons to oxidative stressors (2-4). Increasing evidence points to the contribution of oxidative mechanisms in α-synuclein misfolding/aggregation (5, 6) and stimulation of LRRK2 activity in idiopathic PD (iPD) (7). However, the therapeutic potential of antioxidant approaches for disease modification in PD has never been adequately tested. Agents that have been used in clinical trials, such as α-tocopherol or coenzyme Q_10_, have poor brain penetrance and there has been no good measure of target engagement. Moreover, it may be more efficacious to directly inhibit cellular pro-oxidant systems rather than quenching their final products, reactive oxygen species (ROS).

Although mitochondrial metabolism is considered an important source of ROS that relates to high levels of aerobic respiration in the brain, there are other major sources of ROS that may contribute to redox disturbances in neurodegeneration. NADPH oxidases (NOXs) are electron transferring membrane protein complexes that function as major enzymatic generators of superoxide anion (O_2_^−^) and hydrogen peroxide (H_2_O_2_) (8). Despite the ubiquitous expression of the NOX isoforms 1, 2 and 4 in the central nervous system (9-12), NOX2 is the predominant isoform in brain (13) and has been considered a key player in brain aging (14). While a potential role of NOX2 has been examined in iPD (15) and in PD models (16-19), previous studies have focused on the relevance of NOX2 in PD-related microglial activation. The role of *neuronal* NOX2 in PD pathogenesis remains to be explored.

The subcellular localizations for most of the NOX isoforms have been identified (20) and may vary by cell type. The compartmentalization required for activation of subcellular redox signaling pathways confers specialized cellular functions to the different NOX isoforms. NOX2 is predominantly associated with plasma membrane and endosomes (21). Endosomal trafficking internalizes and re-localizes the NOX2 complex in intracellular organelles, including the endoplasmic reticulum (ER) through inter-organelle membrane contact sites (MCSs) (22). Thus, NOX2 can be associated with the ER and interact with ER-associated organelles, including mitochondria. In certain cellular systems, NOX2 appears intimately related with mitochondrial function in a redox regulatory pathway recently termed *“ROS-induced ROS production*” (RIRP) (23-25). In this scheme, mitochondrially-generated O_2_^−^ stimulates NOX2 activity and *vice versa*. NOX2 (gp91^*phox*^) is constitutively associated with membrane-bound p22^*phox*^, comprising cytochrome b558. Functional NOX2 requires the association with the cytosolic regulatory subunits p67^*phox*^, p40^*phox*^, Rac1 or Rac2, and critical event that is pivotal to NOX2 activation is the association and interaction between the subunits p47^*phox*^ and NOX2 (26). A peptide-based “assembly inhibitor”, NOX2 docking sequence (Nox2ds)-tat, blocks this interaction and is the first widely useful, isoform-specific inhibitor of NOX2 (27-29). Here we report development of a proximity ligation (PL) assay that measures the association of p47^*phox*^ and NOX2 as a surrogate for NOX2 activity. Using this new assay, we examine the mechanism of NOX2 activation, its role in human PD, including the post-translational modification and oligomerization of α-synuclein and activation of LRRK2 kinase activity, as well as the potential and consequences of using the bridged tetrahydroquinoline, CPP11G, a small molecule with selective NOX2 assembly inhibitory function (30, 31), to block neuronal NOX2 activity in vivo in a model of PD.

## Results

### NOX2 activity assay validation

We developed and validated a novel proximity ligation (PL) assay with excellent anatomical and cellular resolution that can rapidly provide information regarding NOX2 activation state, which is based on the in-situ detection of the p47^*phox*^–NOX2 interaction. Mitochondrial ROS are known to activate NOX2 (23-25), and in this context, we found an increased PL signal for NOX2 activity in HEK-293 cells exposed for 24 hours to a sub-lethal concentration (50 nM) of the mitochondrial complex I inhibitor, rotenone (**Figure 1A,B**). The rotenone-treated cells were simultaneously assayed in parallel for superoxide production using dihydroethidium (DHE) and were found to have an increased signal consistent with increased NOX2 activity (**Figure 1A,C**). Using the DHE imaging parameters needed to measure NOX2 activity without saturating the signal, the DHE response to rotenone alone, without NOX2 activation, was undetectable. Rotenone treatment of NOX2^-/-^ HEK-293 cells did not produce a p47^*phox*^–NOX2 PL or DHE signal and, similarly, the NOX2 peptidic assembly inhibitor, Nox2ds-tat (10μM), blocked the interaction and prevented the PL signal – and there was no detectable oxidative response by DHE. The scrambled control peptide variant of Nox2ds-tat was without effect and did not prevent the PL or DHE signals (**Figure 1A-C**).

**Fig. 1.**
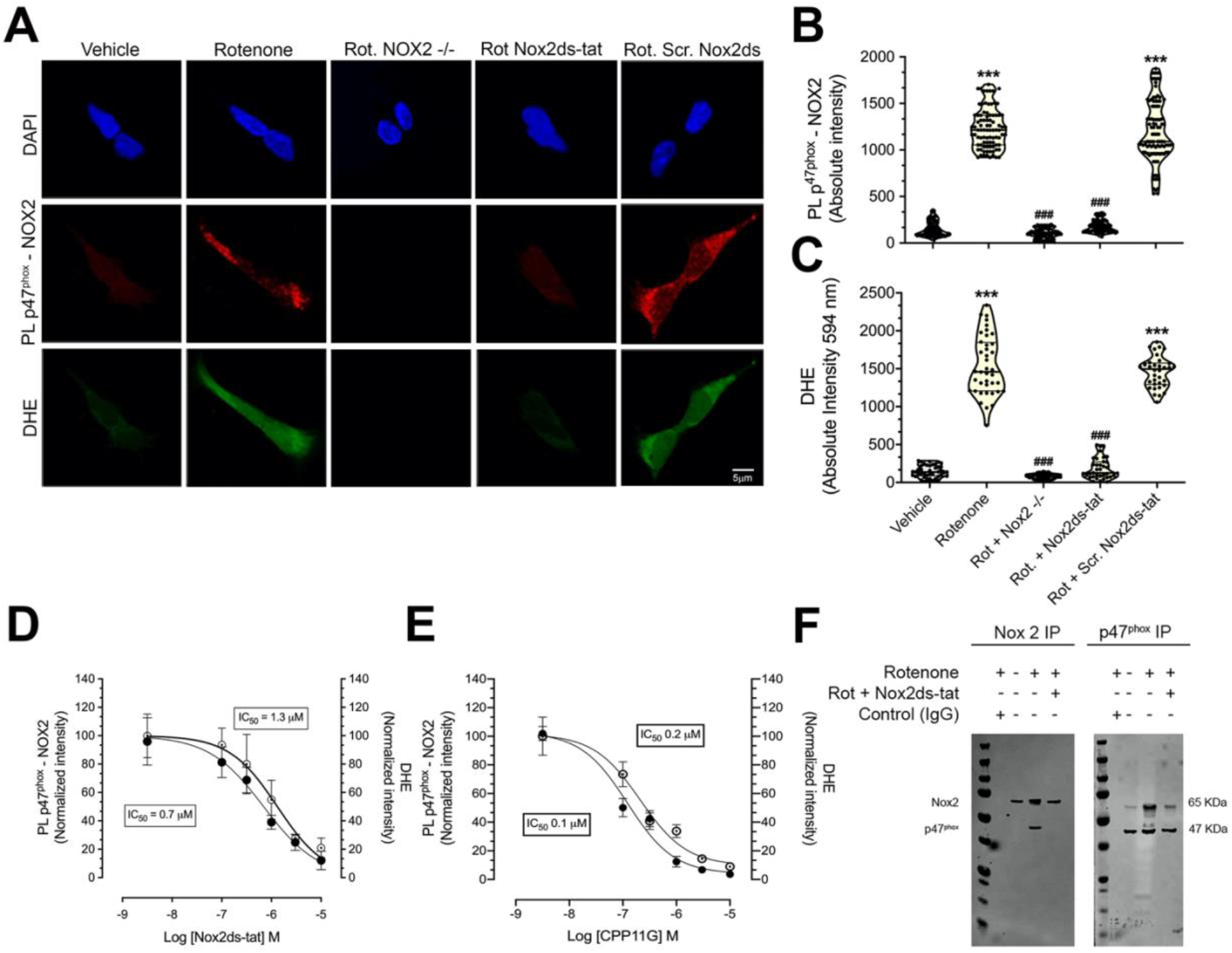
Validation of the p47^phox^-NOX2 proximity ligation assay as an index of activity in situ. **(A**) p47^*phox*^-NOX2 proximity ligation signal (red) and DHE-related fluorescent signal (green) in HEK293 cell lines. Wild-type HEK-293 cells treated with a sub-lethal concentration (50nM) of the complex I inhibitor, rotenone, show increased PL signal for p47^*phox*^-NOX2 interaction and increased cytosolic superoxide production detected by DHE fluorescence. This cellular response to rotenone exposure was prevented in NOX2^-/-^ HEK293 cells. In wild-type cells, co-treatment with Nox2ds-tat, but not by the scrambled variant of Nox2ds-tat peptide prevented the rotenone induced increase in p47^*phox*^-NOX2 and DHE fluorescence. (**B, C**) Quantification of the PL p47^*phox*^-NOX2 signal and DHE-related fluorescence intensity (one-way ANOVA with Bartlett’s post hoc test; 3 independent experiments). (**D, E**) Dose response to NOX2 inhibitors. HEK293 cells were treated with rotenone to activate NOX2 in the absence or presence of increasing concentrations of inhibitor. Quantification of the PL and DHE signals shows that, in a dose-dependent manner, the highly specific NOX2 inhibitors, Nox2ds-tat and CPP11G reduced the PL p47^*phox*^-NOX2 signal (black symbols) (IC_50_: 0.7 µM for Nox2ds-tat and 0.1 µM for CPP11G) paralleled by attenuation of superoxide products (open symbols) (IC_50_: 1.3 µM for Nox2ds-tat and 0.2 µM for CPP11G; n = 3 independent experiments). (**F**) Co-immunoprecipitation assay for NOX2 and p47^*phox*^. In HEK293 cells exposed to 50 nM rotenone for 24 hours, NOX2 and p47^*phox*^ co-immunoprecipitated, suggesting NOX2 activation. Co-treatment with Nox2ds-tat prevented the p47^*phox*^-NOX2 interaction (*i*.*e*., no co-IP) Blot is representative of 3 independent experiments. *** denotes p <0.0001 significance compared vehicle; ### denotes p <0.0001 significance compared to rotenone.

Using rotenone-treated HEK-293 cells, we performed simultaneous dose-response studies of the p47^*phox*^–NOX2 PL and superoxide products (DHE) in response to both the peptidic and small molecule NOX2 assembly inhibitors, Nox2ds-tat and CPP11G (30, 31), respectively (**Figure 1 D,E**). Both inhibitors dose-dependently blocked NOX2 activity (PL signal) and, in parallel, reduced the generation of superoxide (DHE signal). A more conventional NOX2 assay, that lacks anatomical or subcellular resolution, depends on co-immunoprecipitation of p47^*phox*^ with NOX2 to show activation of the enzyme. Using this assay (**Figure 1F**), we confirmed that rotenone treatment activates NOX2 and Nox2ds-tat blocks this effect, as indicated by our PL assay. Thus, we have developed a new PL assay to detect activation of NOX2 that correlates with superoxide production, and which responds in appropriate dose-responsive fashion to structurally distinct classes of specific NOX2 assembly inhibitors. The assay was further validated using gene-edited cells, as well as conventional co-immunoprecipitation approaches.

### Direct evidence of NOX2 activity in human PD and animal models thereof

To determine whether NOX2 activation in nigrostriatal dopamine neurons constitutes a relevant event in the pathogenesis of idiopathic PD, we performed proximity ligation assays (p47^*phox*^– NOX2) in blinded postmortem substantia nigra sections from individuals with PD and from control subjects. Compared to controls, nigrostriatal dopamine neurons from PD cases had a strong proximity ligation signal, indicating NOX2 activation (**Figure 2 A,B;** P < 0.0001 compared to control patients; unpaired t-test). To our knowledge, this is the first demonstration that NOX2 is aberrantly active in nigrostriatal dopamine neurons in individuals with PD.

**Fig. 2.**
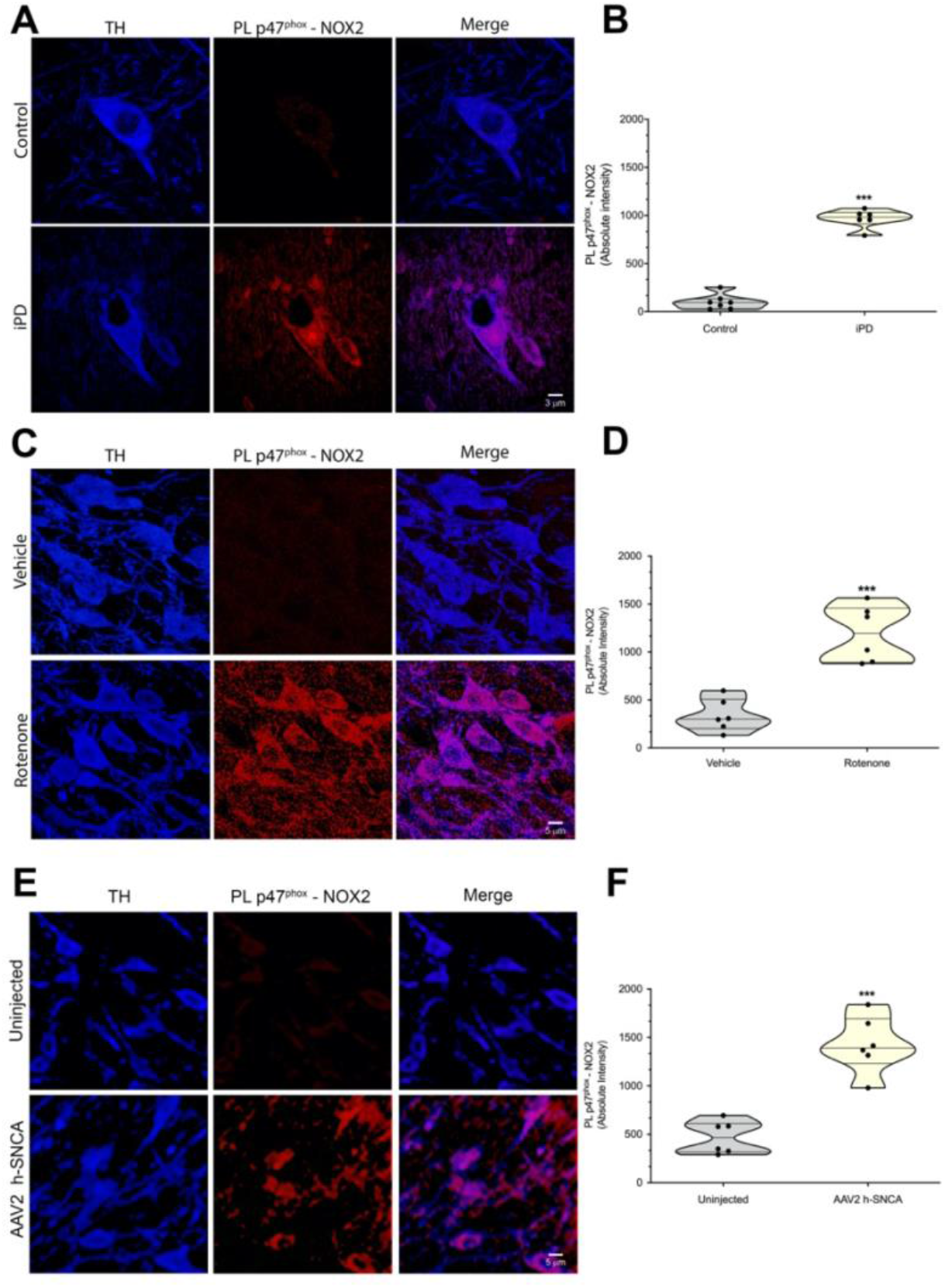
NOX2 activity in nigrostriatal dopamine neurons in human PD postmortem brain tissue and in 2 rat models of PD. (**A**) Representative images of p47^*phox*^-NOX2 proximity ligation signal (red) in midbrain sections from a healthy, age-matched control human brain (top row) and a brain from an individual with iPD (bottom row). Compared to control brains, the PD brains showed a strong p47^*phox*^-NOX2 PL signal in tyrosine hydroxylase (TH) positive neurons (blue). (**B**) Quantification of p47^*phox*^-NOX2 proximity ligation signal in 7 control brains and 6 PD brains (10 – 15 neurons imaged per brain slice). Statistical comparison by unpaired two-tailed *t* test. (**C**) p47^*phox*^-NOX2 proximity ligation signal in the substantia nigra of the brains of rats treated with vehicle (top row) or rotenone (bottom row). In rotenone-treated rats, there was increased p47^*phox*^-NOX2 PL signal, indicating NOX2 activation in nigrostriatal dopamine neurons (TH, blue). (**D**) Quantification of p47^*phox*^-NOX2 proximity ligation signal in nigrostriatal dopamine neurons from control (vehicle) and rotenone-treated rats. Symbols represent individual animals. Statistical comparison by unpaired two-tailed *t* test. (**E**) Shown are p47^*phox*^-NOX2 proximity ligation signal in the substantia nigra from rats that received a unilateral nigral injection of AAV2-*hSNCA*. In the AAV2-*hSNCA*-injected (bottom row), compared to the uninjected hemisphere (top row), a significant increase in p47^*phox*^-NOX2 fluorescent PL signal was observed, indicating that α-synuclein overexpression caused NOX2 activation in nigrostriatal dopamine neurons (TH, blue). (**F**) Quantification of p47^*phox*^-NOX2 proximity ligation signal in nigrostriatal dopamine neurons from the control and AAV-*hSNCA*–injected rat brain hemispheres. Symbols represent mean values from each hemisphere. Statistical comparison by paired two-tailed *t* test. *** denotes p <0.0001 significance compared control, vehicle or uninjected emisphere.

Systemic inhibition of mitochondrial complex I with rotenone reproduces many features of PD in rats (32). Using our proximity ligation assay, we found that, compared to vehicle treatment, the p47^*phox*^–NOX2 interaction signal in dopamine neurons of substantia nigra pars compacta (SNpc) was significantly increased in rotenone-treated rats at endpoint (**Figure 2 C,D;** P < 0.0001 compared to vehicle; unpaired t-test). Similarly, in the rat PD model of AAV2-mediated human α-synuclein over-expression in nigrostriatal neurons (33), we also detected robust NOX2 activity levels in dopamine neurons of the viral injected SNpc, but not in the contralateral non-injected SNpc (**Figure 2 E,F;** P < 0.0001 compared to non-injected side; unpaired t-test). Together, these results indicate that NOX2 activation occurs in SNpc dopamine neurons, but whether it is an early or late phenomenon is unknown to this point.

### Neuronal NOX2 activation is an early phenomenon in the rotenone model

Previous experimental evidence indicates that NOX2 is highly expressed in activated microglial cells in PD patients and in PD animal models (15, 17, 18, 34). The abundance of NOX2 in microglia raises the possibility that the contribution of NOX2 activity to PD pathogenesis may be primarily exerted through microglial oxidative stress-related neuroinflammatory responses. Whether NOX2 activation occurs first in microglia or neurons is also unknown. Using the p47^*phox*^– NOX2 PL assay, we investigated the timing of NOX2 activation in microglia versus nigrostriatal dopamine neurons during the induction of nigrostriatal damage by rotenone *in vivo* in rats (**Figure 3**). Acute administration (1 day) of rotenone was able to induce NOX2 activation in rat dopamine neurons (P < 0.0001 compared to vehicle; unpaired t-test) but not in microglia. Under subacute conditions (rotenone treatment for 5 days), we still observed a PL signal in dopamine neurons (P < 0.0001 compared to vehicle; unpaired t-test) and, at this longer treatment timepoint, there was now activation of microglial NOX2 (P < 0.0001 compared to vehicle; unpaired t-test). Therefore, in this model at least, it appears that NOX2 activation in dopamine neurons constitutes an early event in pathogenesis, whereas microglial NOX2 activity may represent a secondary neuroinflammatory response that supports the progression of the disease.

**Fig. 3.**
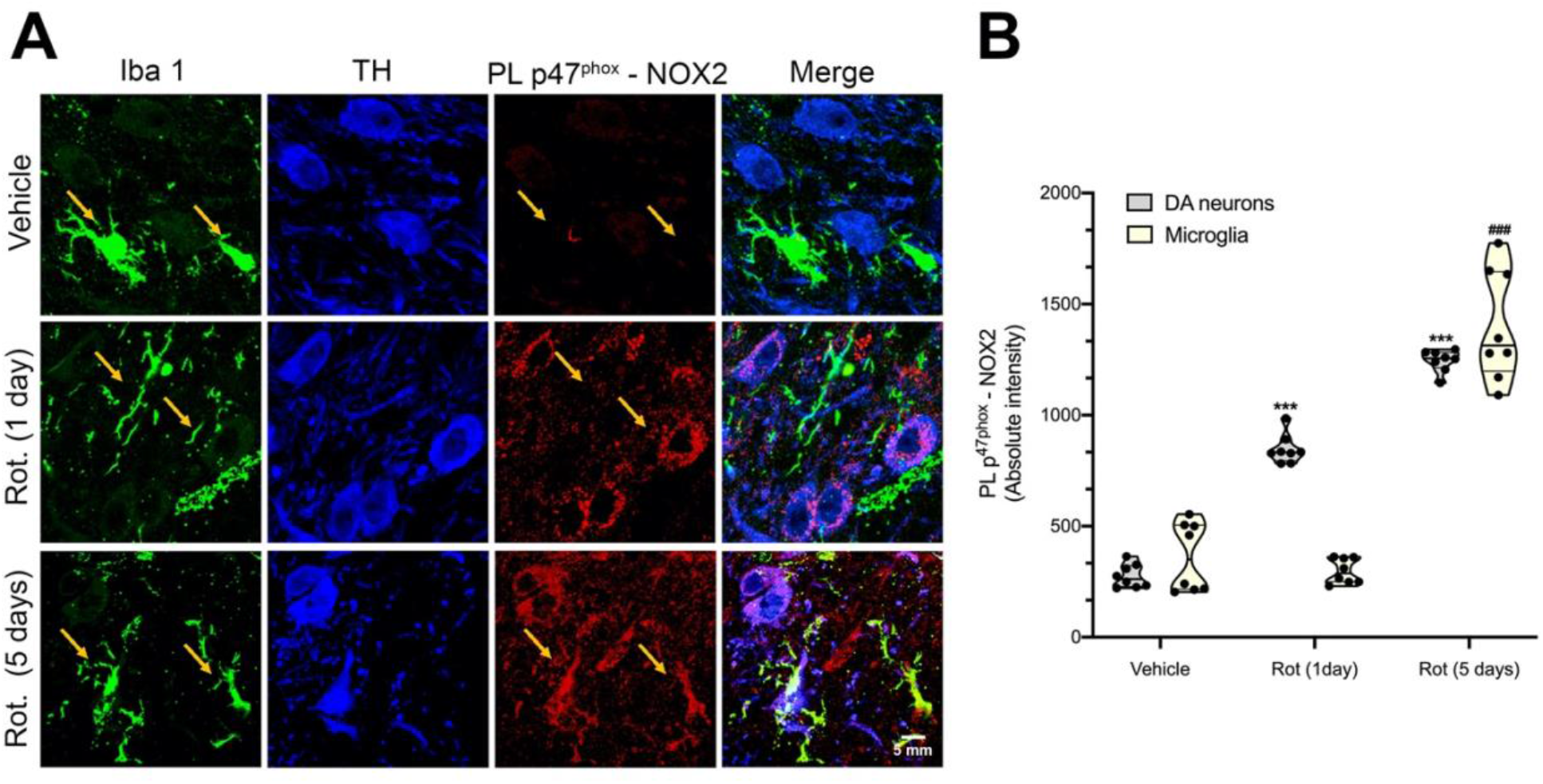
Dopamine neurons show earlier NOX2 activation than microglia in the rotenone model. (**A**) Representative images showing p47^*phox*^-NOX2 PL signal (red) in dopamine neurons (TH, blue) and microglia (Iba-1, green) in substantia nigra sections from rats treated with vehicle (top row) or rotenone for 1 day (middle row) or rotenone for 5 days. In 1-day-treated rats, there was activation of NOX2 in neurons but not microglia. Five days of rotenone treatment elicited an increase in p47^*phox*^-NOX2 proximity signal in both dopamine neurons and in microglia, suggesting that neuronal NOX2 activation and damage might lead to microglial NOX2 activation. (**B**) Quantification of microglial and neuronal NOX2 activity by PL p47^*phox*^-NOX2 signal. Symbols represent single animals. Statistical comparison by one-way ANOVA with post hoc Bartlett’s test. ** denotes p <0.0005 significance compared vehicle *** denotes p <0.0001 significance compared vehicle; ### p <0.0001 significance compared to rotenone.

### Rotenone activation of NOX2 depends on mitochondrial ROS

A growing body of evidence suggests that NOX2 may be activated by mitochondrially-generated ROS in a process known as ROS-induced ROS production (RIRP). To assess this possibility, we employed a simplified model of rotenone treatment of HEK-293 cells to elucidate factors that activate NOX2. Treatment of HEK-293 cells with rotenone (50 nM) for 24 hours elicited NOX2 activation (**Figure 4**), which was prevented by co-treatment with the mitochondrial superoxide scavenger, mito-TEMPO (25 nM), indicating that mitochondrial superoxide production constitutes a critical event leading to NOX2 activation. This confirms a functional link between mitochondrial function and NOX2 activity and also rules out a direct effect of rotenone on NOX2 activity.

**Fig. 4.**
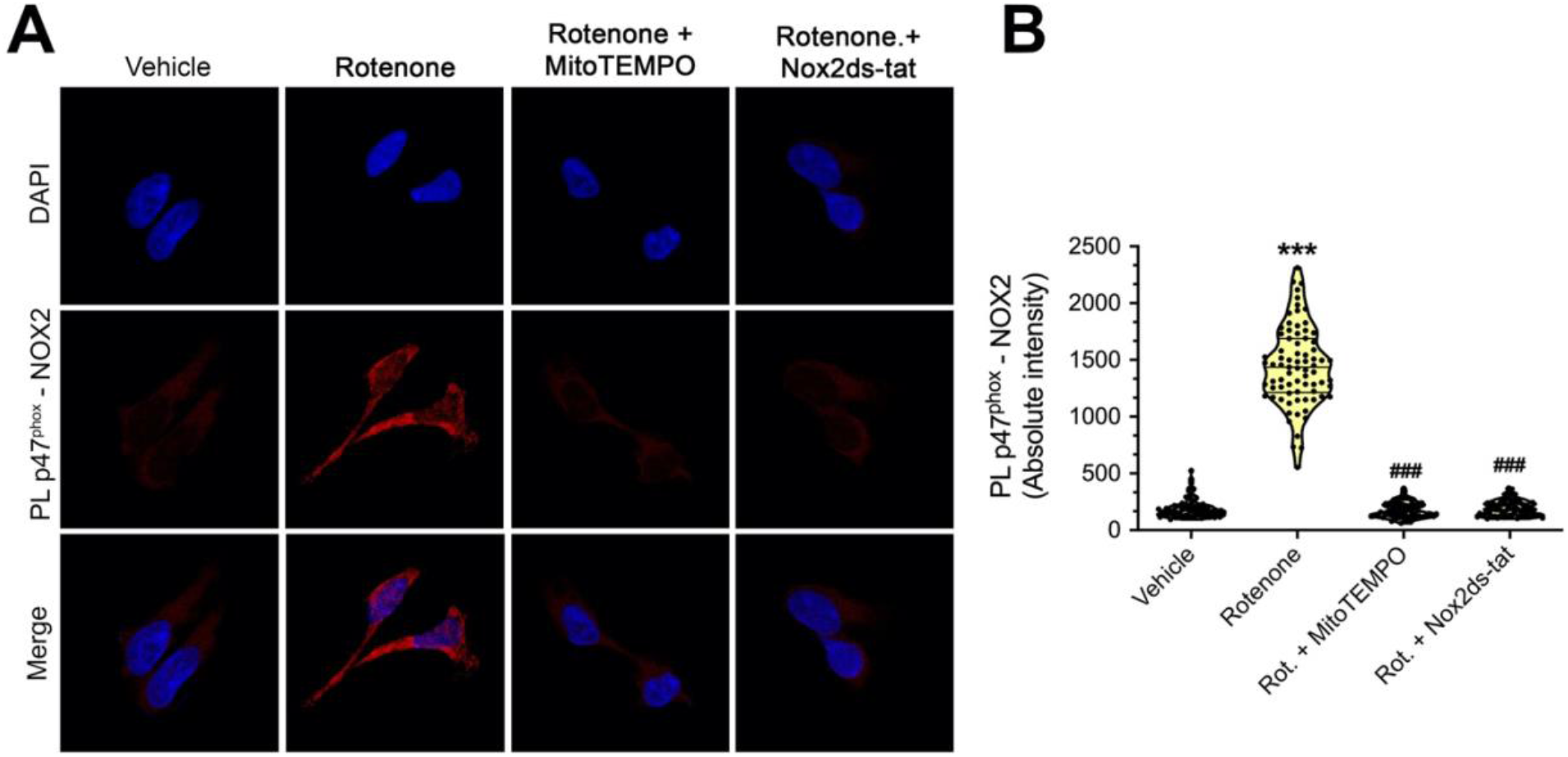
Rotenone-induced mitochondrial superoxide elicits NOX2 activation. (**A**) p47^*phox*^-NOX2 PL signal (red) in wild-type HEK293 cells exposed for 24 hours to vehicle (1^st^ column), rotenone (2^nd^ column), or rotenone plus Mito-TEMPO (3^rd^ column) or rotenone plus Nox2ds-tat (4^th^ column). Treatment with rotenone induced NOX2 activity detected by p47^p*hox*^-NOX2 PL signal (second row). Co-treatment with the mitochondrial superoxide scavenger, mito-TEMPO, prevented rotenone-induced NOX2 activation, consistent with ROS-induced ROS production (RIRP). The NOX2 assembly inhibitor, Nox2ds-tat, prevented rotenone-mediated p47^*phox*^-NOX2 proximity ligation signal, providing further evidence on the specificity of the assay. (**B**) Quantification of PL signal for NOX2 activity by pL P47^*phox*^-NOX2 signal. Symbols represent the average of single cell fluorescence intensity analyzed in one 100X microscopic field. Statistical comparison by one-way ANOVA with post hoc Tukey’s test on three independent experiments. *** denotes p <0.0001 significance compared vehicle; ### p <0.0001 significance compared to rotenone.

### Effects of NOX2 activity on α-synuclein

Post-translational modifications (PTM) of α-synuclein are believed to be important in the pathogenesis of PD and the associated SNpc dopamine neuron degeneration (35). Previous evidence from our laboratory defined critical roles of oligomeric α-synuclein and phospho-serine129-α-synuclein (pSer129-Syn) in mitochondrial protein import impairment (6). However, how the formation of certain neurotoxic PTMs of α-synuclein occurs is still unclear. Oxidative stress-related PTMs of α-synuclein, including 4-hydroxy-2-nonenal-α-synuclein (HNE-αSyn) and nitrated α-synuclein (NO-αSyn) have been shown to promote α-synuclein oligomerization (36). Here, we found that treatment of HEK-293 cells with rotenone for 24 hours increased levels of total α-synuclein (**Figure 5 A, B**) and pSer129-Syn (**Figure 5C, D**). Notably, the effects of rotenone were prevented in NOX2^-/-^ HEK-293 cells (P < 0.0001 compared to wild-type-treated cells; ANOVA with Bartlett’s correction) and by Nox2ds-tat co-treatment (P < 0.0006 compared to wild-type-treated cells; ANOVA with Bartlett’s correction).

**Fig 5.**
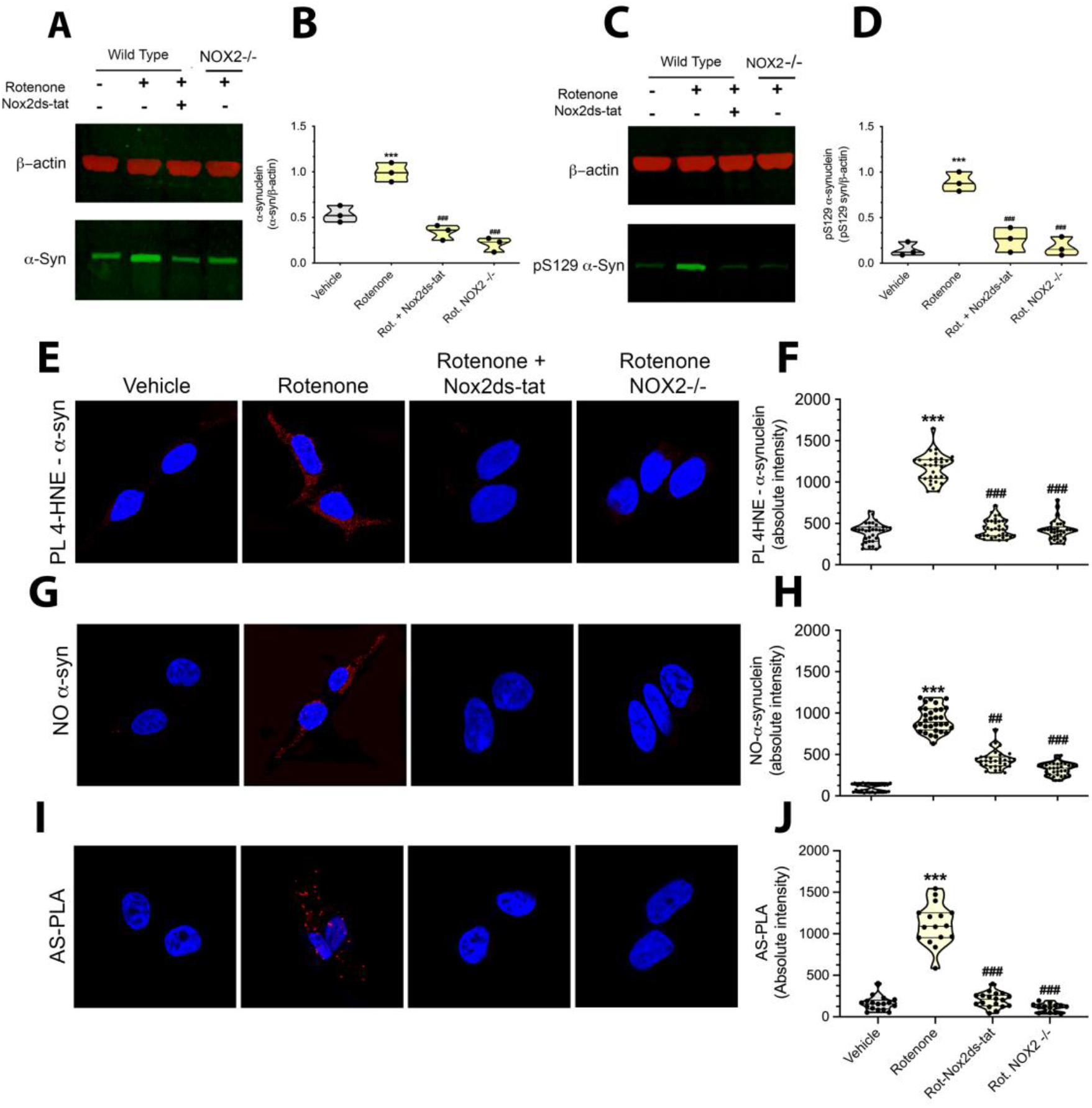
NOX2-dependent accumulation and formation of PTM forms of α-synuclein. (**A, C**) Western blot analysis for α-syn and pS129-α-syn in wild-type cells and CRISPR/Cas9 NOX2^-/-^ HEK293 cells (NOX2^-/-^). After 24 hours of rotenone exposure, wild-type cells showed an accumulation of (A) total α-syn and (C) pS129-α-syn. NOX2^-/-^ cells treated with rotenone showed no increase in total α-synuclein accumulation and phosphorylation. The same effect was observed in wild-type cells co-treated with Nox2ds-tat. (**B**) Quantification of total α-syn and (**D**) pS129-α-syn. Symbols represent a single replicate. Statistical comparison by one-way ANOVA with post hoc Bartlett’s test. Confocal analysis in wild-type and NOX2^-/-^ cells for (**E**) 4-HNE-α-syn (detected as 4-HNE–α-syn PL signal), (**G**) NO-α-syn (by immunocytochemistry) and (**I**) oligomeric α-syn (as AS-PLA). Cells exposed to rotenone for 24 hours showed significant increase of 4-HNE–α-syn proximity signal (**E**) and immunoreactivity for NO-syn (**I**). Delayed formation of oligomeric α-syn was also observed after 48 hours of rotenone exposure (**G**). These PTM forms of α-synuclein were not observed in NOX2^-/-^ treated with rotenone or wild-type cells co-treated with Nox2ds-tat. (**F**) Quantification of PL 4-HNE–α-syn signal, (**H**) fluorescence intensity relative to NO-α-syn and (**J**) AS-PLA fluorescence intensity. Symbols represent the average of single cell fluorescence intensity analyzed in one 100X microscopic field. Statistical comparison by one-way ANOVA with post hoc Bartlett’s test on three independent experiments. *** denotes p <0.0001 significance compared vehicle; ### p <0.0001 significance compared to rotenone.

We further investigated the role of NOX2 activity in the formation of PTM forms of α-synuclein, including HNE-αSyn, NO-αSyn and oligomeric α-syn in rotenone-exposed HEK-293 cells. The PL assay for HNE-αSyn adducts and the immunofluorescence analysis for NO-αSyn revealed a robust increase of cellular levels of these oxidative stress-modified forms of α-synuclein in rotenone-treated HEK-293 cells. NOX2 knockout or Nox2ds-tat co-treatment prevented both rotenone-induced HNE-αSyn adducts and NO-αSyn (**Figure 5 E, F** for HNE-αSyn; P < 0.0001 compared to wild-type-treated cells; ANOVA with Bartlett’s correction. **Figure 5 G, H** for NO-αSyn; P < 0.0001 compared to wild-type-treated cells; ANOVA with Bartlett’s correction). We also assayed the formation of oligomeric α-synuclein by proximity ligation assay (AS-PLA; as described(37)). Treatment with rotenone for 48 hours elicits the formation of α-synuclein oligomers (**Figure 5 I & J**) and this was prevented in gene-edited NOX2^-/-^ HEK-293 cells (p < 0.0001 compared to wild-type-treated cells; ANOVA with Bartlett’s correction) and by Nox2ds-tat co-treatment in wildtype HEK-293 cells (p < 0.0001 compared to wild-type-treated cells; ANOVA with Bartlett’s correction). Together these results suggest that NOX2 mediates formation of HNE-αSyn and NO-αSyn species and is responsible for α-synuclein oligomerization.

Neurons and microglia co-express other NOXs isoforms, among which the most abundant are NOX1 and NOX4. We developed CRISPR/Cas9 gene-edited NOX1^-/-^, NOX2^-/-^ and NOX4^-/-^ HEK-293 cells (**Supplemental Figure 1 A, B**) to investigate the role of these NOXs isoforms in rotenone-mediated formation of HNE-α-syn. Only knockout of NOX2 (not NOX1 or NOX4) prevented rotenone-induced formation of HNE-α-syn, implicating for the first time NOX2 activity in the formation and accumulation of cytotoxic forms of α-synuclein.

We previously reported that certain modified forms of α-synuclein, including pSer129-α-syn and oligomeric α-syn, bind to TOM20 of the mitochondrial protein import machinery and cause mitochondrial protein import impairment and mitochondrial dysfunction (6). In the current study, we assessed the downstream consequences of NOX2-induced α-synuclein PTMs. Proximity ligation assays (α-synuclein–TOM20) in HEK-293 cells exposed to rotenone for 24 hours revealed a strong α-synuclein–TOM20 interaction and a corresponding diffuse cytoplasmic redistribution of Ndufs3 (an imported nuclear-encoded Complex I subunit), indicative of impaired mitochondrial protein import. Exposure to Nox2ds-tat prevented the α-synuclein–TOM20 interaction (**Supplemental Figure 2**; P < 0.0001; unpaired T-test), suggesting that oxidative modifications of α-synuclein may constitute a link between NOX2 activation and downstream mitochondrial dysfunction.

### A brain-penetrant inhibitor prevents nigrostriatal NOX2 activation and its downstream effects

To provide evidence of CPP11G blood brain barrier (BBB) permeability and persistence in brain tissue, we performed mass spectrometry analysis to measure the presence of CPP11G in rat brains 24 hours after a single oral administration of CPP11G (15 mg/kg). At this prolonged timepoint, CPP11G was readily detectable in whole brain lysates, indicating BBB penetrance and persistence of the NOX2 inhibitor (**Supplemental Figure 3A**). Similarly, mass spectrometry analysis after a single intraperitoneal dose of rotenone (2.8 mg/Kg), revealed its residual presence in brain 24 hours after injection (**Supplemental Figure 3B**). Mass spectrometry also showed that when rotenone was co-administered with CPP11G, it did not alter the brain concentration of the NOX2 inhibitor. Moreover, co-administration of CPP11G did not change the brain concentration of rotenone (**Supplemental Figure 3A,B**), suggesting that the two compounds do not interfere with the metabolism of each other.

In this study, we also examined whether the NOX2 inhibitor, CPP11G, was able to block rotenone induced NOX2 activation in SNpc dopamine neurons *in vivo*, as well as its downstream sequelae. Twenty-four hours after a *single* rotenone administration (2.8 mg/kg; i.p.), nigrostriatal neurons showed a marked elevation of NOX2 activity, detected as PL signal for p47^*phox*^–NOX2 (**Supplemental Figure 3C,D**). When CPP11G was co-administered with rotenone, it prevented the rotenone induced NOX2 activation in dopamine neurons when assessed 24 hours after treatment.

The rotenone model of PD reproduces many features of the human disease, including oxidative stress, LRRK2 activation, and reduced mitochondrial protein import (2, 3, 6, 7). To determine whether a NOX2 inhibitor could block more prolonged rotenone-induced NOX2 activation, and to assess PD-related downstream effects, we treated rats for 5 days with rotenone (2.8 mg/kg/day, i.p.) with or without CPP11G (15 mg/kg/day, p.o.). Daily systemic administration of rotenone causes a progressive weight loss in rats, likely due to its deleterious effects on mitochondria. Here, 5 days of rotenone treatment caused a 10% weight loss compared to vehicle control animals. Co-treatment with CPP11G prevented rotenone-induced loss of weight, suggesting a critical role of NOX2 in the systemic toxicity of rotenone exposure (**Supplemental Figure 4**).

When assayed 24 hours after cessation, 5 days of rotenone treatment was found to have induced a robust increase in p47^*phox*^–NOX2 proximity ligation signal in nigrostriatal dopamine neurons, which was associated with an increase in lipid peroxidation, detected as 4-HNE levels (**Figure 6**). Cotreatment with CPP11G effectively blocked the rotenone-induced activation of NOX2 (P < 0.0001, two-way ANOVA with Bartlett’s correction) and formation of 4-HNE (P < 0.0001; two-way ANOVA with Bartlett’s correction).

**Fig. 6.**
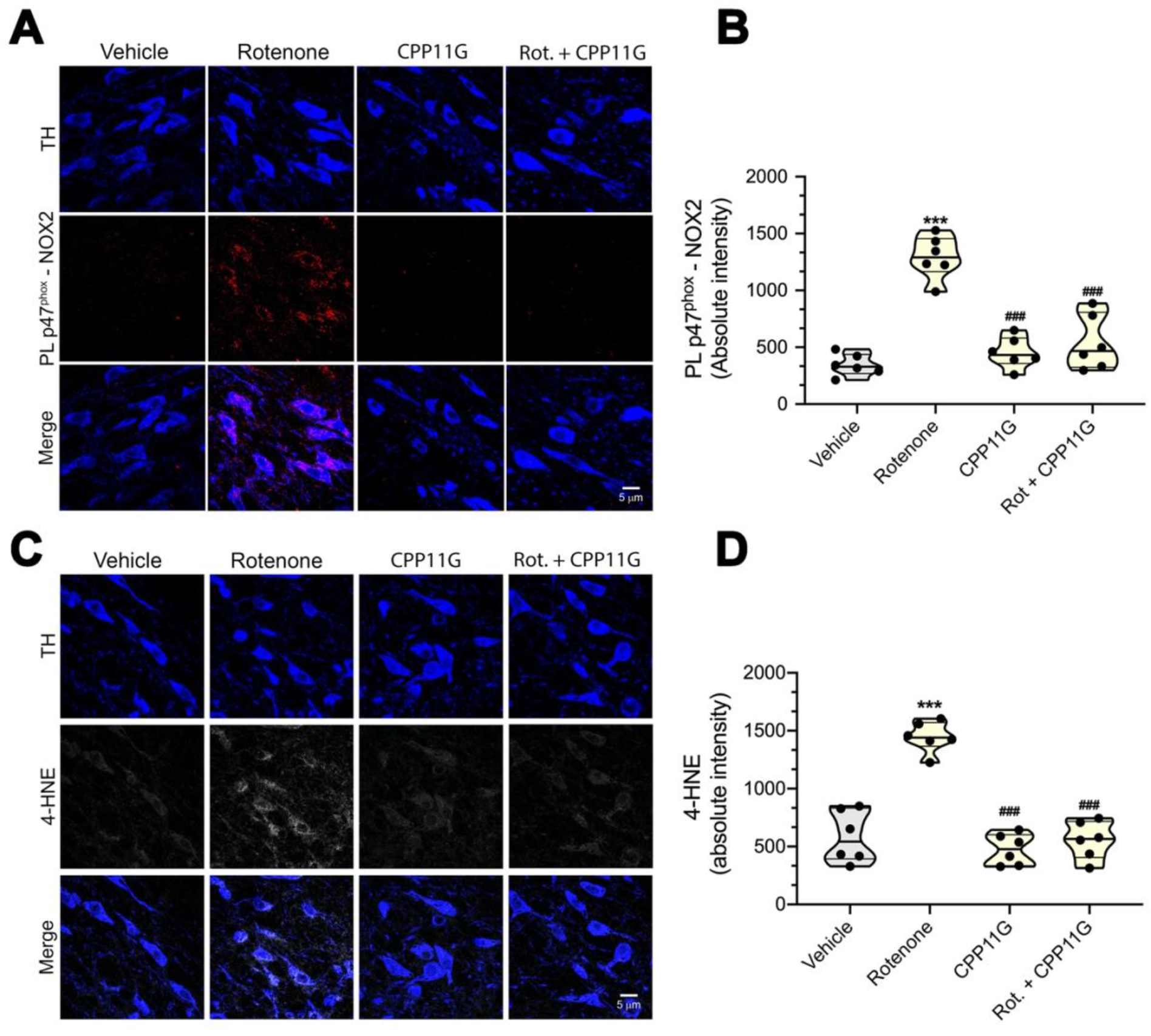
In vivo nigrostriatal NOX2 activation and oxidative damage in the rotenone model of PD are prevented by a brain penetrant NOX2 inhibitor. (**A**) Representative images of p47^*phox*^-NOX2 PL signal (red) detecting the interaction of p47^*phox*^ with NOX2. After 5 days of rotenone treatment, nigrostriatal dopamine neurons (TH, Blue) showed a sustained NOX2 activity that was prevented in animals that received co-treatment with oral CPP11G. (**B**) Quantification of p47^*phox*^-NOX2 proximity ligation signal. Each symbol represents an individual animal. Comparison by ANOVA with post hoc Bonferroni correction. (**C**) Fluorescence images of 4-hydroxynonenal (4-HNE) immunohistochemistry (gray) as marker of lipid oxidation. The analysis was performed in nigrostriatal dopamine neurons (TH; blue) from rat brains. Five days of rotenone treatment caused oxidative stress and a significant increase of 4-HNE in nigrostriatal dopamine neurons. Co-treatment with CPP11G prevented lipid peroxidation. (**D**) Quantification of 4-HNE fluorescence signal. Each symbol represents an individual animal. Comparison by ANOVA with post hoc Bartlett’s correction. *** denotes p <0.0001 significance compared vehicle; ### p <0.0001 significance compared to rotenone.

We previously reported that LRRK2 kinase activity can be stimulated by oxidative mechanisms, which may involve NOX2 activity (7). Here, we found that rats treated with rotenone for 5 days also showed robust LRRK2 kinase activity in dopamine neurons, and this was blocked by co-treatment with CPP11G (**Figure 7A,B**), indicating LRRK2 kinase activity is a downstream effect of NOX2 activation in an epidemiologically relevant model of PD.

**Fig. 7.**
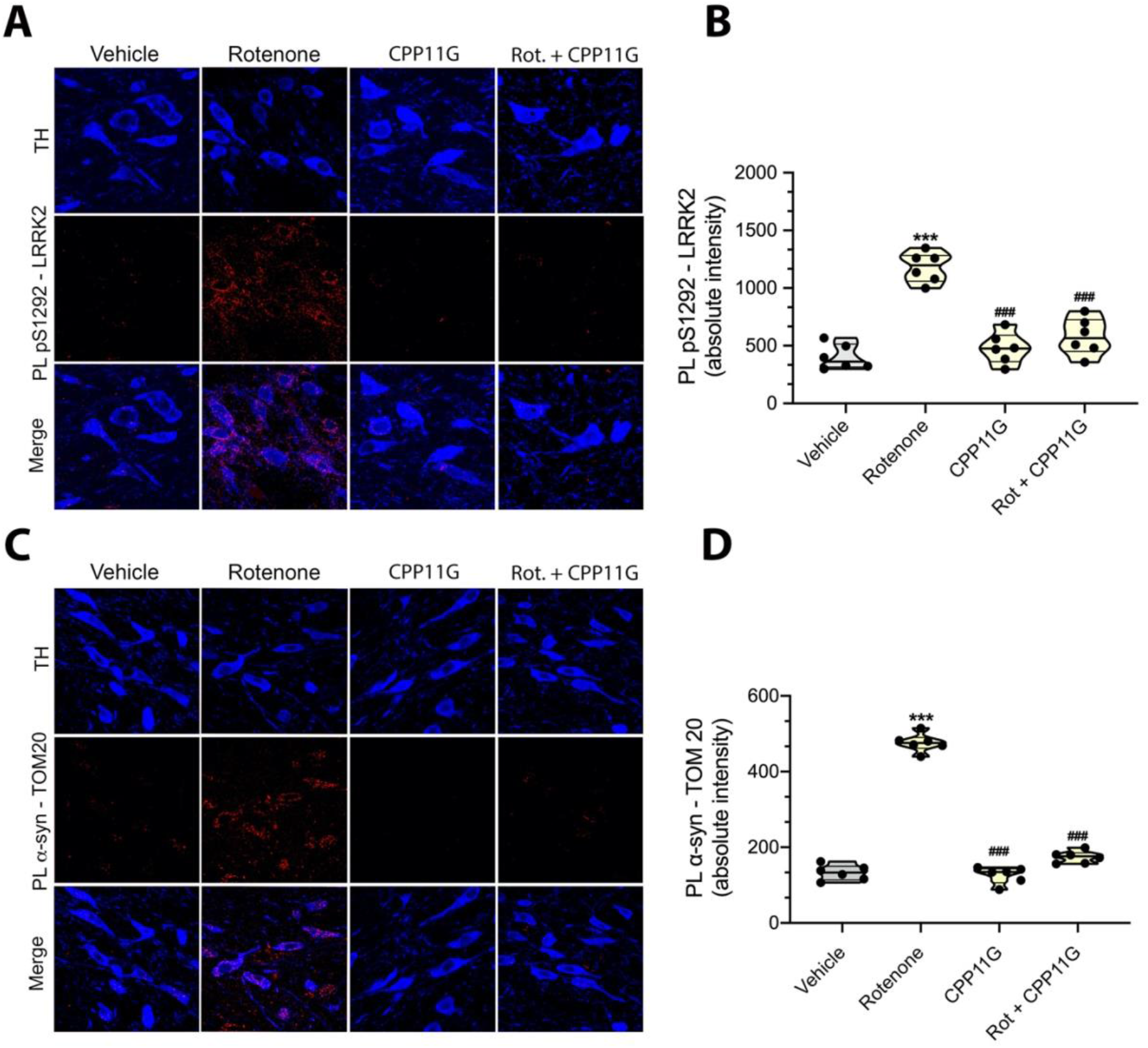
The brain penetrant NOX2 inhibitor, CPP11G, prevents rotenone-induced LRRK2 activation and α-synuclein-TOM20 interaction in rat nigrostriatal dopamine neurons. (**A**) Shown are the pSer1292-LRRK2 PL signal (red) in rats treated with vehicle (1^st^ column), rotenone alone (2^nd^ column), CPP11G alone (3^rd^ column), or rotenone + CPP11G (4^th^ column). After 5 days of treatment, rotenone elicited a significant increase in pSer1292-LRRK2 signal, indicating LRRK2 kinase activation in nigrostriatal dopamine neurons. When co-administrated with rotenone, CPP11G was effective in blocking rotenone mediated LRRK2 kinase activity. (**B**) Quantification of pSer1292 proximity ligation signal in nigrostriatal dopamine neurons. Symbols represent individual animals. Comparison by ANOVA with Bartlett’s correction. (**C**) PL assay detects interaction between α-synuclein and the receptor of the mitochondrial outer membrane protein import machinery, TOM20 (red) in rat. Rotenone treatment induced a strong PL signal for α-synuclein-TOM20 interaction, suggesting impaired mitochondrial protein import. Co-administration of CPP11G prevents rotenone-mediated α-synuclein-TOM20 PL signal. (**D**) Quantification of α-synuclein-TOM20 PL signal in nigrostriatal dopamine neurons. Symbols represent individual animals. Comparison by ANOVA with post hoc Bonferroni correction. *** denotes p <0.0001 significance compared vehicle; ### p <0.0001 significance compared to rotenone.

Similar to our previous results (6) in rats that were treated with rotenone to endpoint (10 – 14 days), here, when rats were treated with rotenone for 5 days, we observed a strong α-syn–TOM20 proximity ligation signal (P < 0.0001, two-way ANOVA with Bartlett’s correction). Cotreatment with CPP11G prevented the interaction of α-synuclein with TOM20 (**Figure 7C,D**). Thus, both the accumulation of PTM forms of α-synuclein (**Figure 5**) and its toxic consequences appear to be downstream of NOX2 activity.

## Discussion

Results of the current study validate a new *in situ* assay for NOX2 and show that substantia nigra neuronal NOX2 is activated in idiopathic PD. *In vitro* and *in vivo* experiments showed that NOX2 is activated by mitochondrial ROS and that, in turn, NOX2 activity is responsible for post-translational modification and oligomerization of α-synuclein and activation of wildtype LRRK2 in a model of PD. Treatment of rats with a brain-penetrant, specific inhibitor of NOX2 prevented NOX2 activation *in vivo* and blocked its downstream effects on α-synuclein and LRRK2.

Until now, it has been difficult or impossible to assess the activity of NOX2 *in situ*, in fixed tissue, or with a cellular level of resolution. The recruitment of p47^*phox*^ to the NOX2 catalytic membrane complex constitutes the key step in NOX2 activation. In this context, we developed a proximity ligation assay to detect the *in-situ* interaction of p47^*phox*^ with NOX2, which allows assessment of NOX2 activation state with a cellular level of resolution (*e*.*g*., in dopamine neurons vs. surrounding microglia). Mitochondrial ROS are known to activate NOX2 and accordingly, when HEK-293 cells were treated with the mitochondrial complex I inhibitor, rotenone, there was a marked increase in the p47^*phox*^–NOX2 PL signal that correlated with the cytoplasmic oxidative stress, as assessed by DHE. In contrast, in gene-edited NOX2^-/-^ cells, rotenone elicited neither a PL signal nor a detectable increase in DHE signal.

Nox2ds-tat is a peptidic inhibitor that mimics a sequence in the cytosolic B-loop of NOX2 (27, 38). As such, it binds the cytosolic regulatory protein, p47^*phox*^, which in turn prevents its translocation/binding to, and activation of, the NOX2 enzyme complex. This highly specific inhibitor blocked the rotenone-induced p47^phox^–NOX2 PL signal as well as the DHE signal with IC_50_s of 0.7 and 1.3 µM, respectively. Similarly, the small molecule NOX2 inhibitor, CPP11G (30), reduced both the NOX2 PL and DHE signals. Further, the dose-dependency of the assay relative to inhibitor exposure indicates the ability of the PL assay to provide a quantitative measure of target (NOX2) engagement. Finally, we note that the PL assay results are entirely consistent with conventional co-immunoprecipitation assays for NOX2 and p47^*phox*^, further validating the method.

Previous work, using other assays, different tissues and cell types, and other experimental conditions has found that there is cross-talk between mitochondria and NOX isoforms (24, 39, 40). As described above, we found that the mitochondrial complex I inhibitor, rotenone, generated a p47^*phox*^–NOX PL signal, increased O_2_^−^ production, and enhanced association of p47^*phox*^ with gp91^*phox*^, as assessed by co-immunoprecipitation. This rotenone-induced activation of NOX2 was blocked by treatment with mito-TEMPO, a mitochondrially targeted O_2_^−^ scavenger. This indicates that (i) mitochondrially-derived ROS can activate NOX2, and (ii) rotenone does not activate NOX2 directly, as suggested by others (41). Given that mitochondrial impairment is strongly implicated in PD pathogenesis, and that it can activate NOX2, it was logical to examine the activation state of NOX2 in brains of people who died with PD.

Using our p47^*phox*^–NOX2 PL assay, we detected markedly enhanced NOX2 activity in nigrostriatal dopamine neurons in postmortem brain tissue from patients with iPD compared to age-and sex-matched controls. We were able to reproduce this finding in the rotenone and the AAV2-h*SNCA* rat models of PD. Microglial activation (neuroinflammation) is a pathological feature of PD brains (42), and increased NOX2 expression in such post-mortem specimens correlates with microglial activation and proliferation (15). Moreover, in animal models of PD, there is upregulation of NOX2 (gp91^*phox*^) and phosphorylation of p47^*phox*^ in SNpc microglial cells (34, 43). However, neurons express NOX2 as well (10, 12) and it is unclear whether neuronal or microglial NOX2 activation occurs first. In order to delineate the relative temporal profiles of NOX2 activation in microglia versus neurons in SNpc, we treated rats with rotenone for a single day, or for 5 days. Importantly, neither treatment regimen causes detectable neurodegeneration. We found that 1 day of rotenone activated NOX2 in nigral dopamine neurons, but not in surrounding microglia. By 5 days, however, microglia also showed NOX2 activation. This suggests that neuronal NOX2 activation (and cell damage) precedes microglial NOX2 activation and may be more proximate to disease initiation.

The pathological accumulation of toxic forms α-synuclein in dopaminergic neurons is believed to be central to PD-related nigrostriatal neurodegeneration. We have previously reported that rotenone-induced mitochondrial impairment leads to posttranslational modifications (PTMs) of α-synuclein, including oligomers and pSer129-α-syn. In the current study, we found that NOX2 activation is an essential step in the process by which rotenone causes these and other toxic species of α-synuclein to form. Indeed, in NOX2^-/-^ cells, rotenone had no effect on α-synuclein, and in wildtype cells, NOX2 inhibitors blocked rotenone’s effects. This appears to be a specific effect of NOX2, since knockout of NOX1 and NOX4 isoforms had no effect on rotenone-induced PTM of α-synuclein.

Oxidative stress-related PTM forms of α-synuclein, including modification by 4-hydroxy-2-nonenal and nitration, have a higher propensity to form oligomers than unmodified monomeric α-synuclein (44, 45). In this context, HNE-α-syn is particularly and selectively toxic to dopamine neurons (36, 46). We detected NOX2 activity-dependent accumulation of HNE-α-syn and NO-syn 24 hours after rotenone exposure and, given their reported enhanced propensity to aggregate, we assessed the accumulation of α-synuclein oligomers in wild type HEK-293 and NOX2^-/-^ cells exposed to rotenone, using the AS-PLA assay developed by Roberts and colleagues (37). We found that, in contrast to HNE-α-syn and NO-α-syn, 24 hours of rotenone is insufficient to induce a detectable amount of α-synuclein oligomers; however, a strong AS-PLA signal was detected after 48 hours of rotenone exposure. The AS-PLA signal was prevented in NOX2 null cells or by Nox2ds-tat co-treatment in wild-type cells. Thus, it is conceivable that NOX2 activity drives the formation of α-synuclein aggregates via increased levels of HNE-α-syn and NO-α-syn. Moreover, this work provides a compelling link between neuronal NOX2 and α-synuclein pathology and toxicity.

Using a specific, brain-penetrant, small molecule inhibitor of NOX2, CPP11G, we were able to perform *in vivo* studies in a rat model of PD. Treatment of rats with rotenone for 5 days resulted in a marked activation of NOX2 and accumulation of HNE in SNc dopaminergic neurons, both of which were blocked by co-administration of CPP11G. We previously reported that wildtype LRRK2 is activated, via oxidative mechanisms, in idiopathic PD, and this was reproduced in rotenone-treated rats. Here, we found that rotenone induced LRRK2 activation was blocked effectively by inhibiting NOX2 *in vivo* with CPP11G, suggesting that NOX2 activity is a requisite downstream effector of mitochondrial ROS. Given our findings that (i) LRRK2 is activated in nigral neurons in idiopathic PD (7), (ii) NOX2 activity is required for wildtype LRRK2 activation *in vivo*, and (iii) NOX2 is activated in nigral neurons in idiopathic PD, it appears this mechanism is operative in the human disease.

As discussed earlier, NOX2 activity appears to be responsible for the aberrant formation of PTM and oligomeric forms of α-synuclein. NOX2 activity also activates wildtype LRRK2, which leads to autophagic dysfunction and impaired degradation of pathological forms of the protein (7, 47). Thus, NOX2 contributes to both the formation and the compromised disposal of toxic species of α-synuclein. In this context, oligomeric α-synuclein and pSer129-α-syn are among the species of α-synuclein that bind to TOM20 and impair mitochondrial function (6, 7). Our *in vivo* experiments showed that NOX2 inhibition also prevented this form of α-synuclein-induced mitochondrial toxicity.

Together, our results suggest that neuronal NOX2 plays a pivotal role in the pathogenesis of PD (**Supplemental Figure 5**). It is activated by mitochondrial ROS and serves to amplify that signal, producing sufficient O_2_^−^ to cause lipid peroxidation, posttranslational modification and oligomerization of α-synuclein, and activation of LRRK2 kinase activity in nigral dopamine neurons. Further, by both directly facilitating formation of pathogenic species of α-synuclein and indirectly inhibiting their autophagic degradation (via LRRK2 activation), NOX2 activity also amplifies the downstream mitochondrial toxicity of α-synuclein. In light of this apparent feed forward cycle, NOX2 presents an attractive target for the simultaneous therapeutic amelioration of ongoing α-synuclein toxicity and aberrant LRRK2 activity. Our *in vivo* studies, using an epidemiologically relevant model of PD (48), provide strong preclinical support for this strategy and for a selective and potent class of novel inhibitors that target NOX2 in disease.

## Materials and methods

### Study design

This study was designed to assess the role of NOX2 in iPD. To do so, we developed new proximity ligation assays to assess *in-situ* interaction NOX2-p47^*phox*^ as surrogate of NOX2 activity. This assay was validated using CRISPR/Cas9 genome-edited NOX2^-/-^ HEK-293 cells and two highly specific NOX2 inhibitors. The validated assay was then used to assess the NOX2 activation state in iPD postmortem brain tissue and in two rat models of PD. Additional studies examined the role of NOX2 activity as well as downstream consequences of NOX2 activity in vivo.

All *in vitro* experiments were replicated at least three times, and key validation studies identifying CRISPR/Cas9-edited cell lines by means of proximity ligation assays were analyzed by blinded assessors. *In vivo* experiments using rotenone treatment, with or without concomitant CPP11G treatment, were performed in a single cohort of rats (n = 6 per active treatment group), and outcomes were analyzed by blinded assessors. Rats were randomized to the treatment group. There was no exclusion of outliers.

### CRISPR/Cas9 genome editing of HEK-293 cells to produce NOXs knockouts cell lines

NOX1, NOX2 and NOX4 knockouts cell lines were generated by Synthego Corporation using CRISPR/Cas9 genome editing technology. Cell clones were grown and expanded for polymerase chain reaction (PCR) and DNA sequence analyses. To confirm gene editing of the NOXs genes, PCR products were sequenced.

### Proximity ligation assays

Proximity ligation (Duolink; Sigma Aldrich -https://reedd.people.uic.edu/ReedLabPLA.pdf) was performed as previously described (6, 7) in 4% PFA (paraformaldehyde)–fixed tissue or cells. Samples were incubated with specific primary antibodies to the proteins to be detected. Secondary antibodies conjugated with oligonucleotides were added to the reaction and incubated. Ligation solution, consisting of two oligonucleotides and ligase, was added. In this assay, the oligonucleotides hybridize to the two proximity ligation probes and join to a closed loop if they are in close proximity. Amplification solution, consisting of nucleotides and fluorescently labeled oligonucleotides, was added together with polymerase. The oligonucleotide arm of one of the proximity ligation probes acts as a primer for “rolling-circle amplification” using the ligated circle as a template, and this generates a concatemeric product. Fluorescently labeled oligonucleotides hybridize to the rolling circle amplification product. The proximity ligation signal was visible as a distinct fluorescent spot and was analyzed by confocal microscopy. Control experiments included routine immunofluorescence staining of the proteins of interest under identical experimental conditions.

### Statistical analyses

Each result presented here was derived from three to six independent experiments. For simple comparisons of two experimental conditions, two-tailed, unpaired t tests were used. When AAV vector was injected into one hemisphere of the rat brain and the other hemisphere was used as a control, two-tailed paired t tests were used. For comparisons of multiple experimental conditions, one-way or two-way ANOVA was used, and if significant, overall post hoc Bartlett’s or Tukey’s corrections for multiple pairwise comparisons were made. P values less than 0.05 were considered significant.

## Author contributions

M.T.K. designed, performed, and analyzed the proximity ligation experiments and edited the manuscript; E.K.H. was responsible for molecular biology and created and validated cell lines; K.F. performed *in vivo* rotenone experiments; C.R.B. performed and analyzed many of the proximity ligation experiments *in vivo*; M.F. performed mass spectrometry analyses to define CPP11G brain penetrance; A.Z. did *in vivo* α-synuclein overexpression work; S.L.C. performed histology and immunostainings in rat brains and human specimen; J.K.K. provided human neuropathology expertise and samples; E.C.P. and P.J.P. provided pharmacological expertise and Nox2ds-tat and CPP11G; E.A.B. provided input and collaboration on the *in vivo* α-synuclein overexpression experiments; T.G.H. provided guidance and helped to design experiments; J.T.G. provided guidance and helped to design experiments and edited the manuscript; R.D.M. supervised the project, designed and analyzed the experiments, and wrote the paper.

## Data and materials availability

All data associated with this study can be found in the paper and the Supplementary Materials.

## Supplementary Materials

### Materials and Methods

Fig. S1. NOX2 is the key NADPH oxidase isoform in the induction of PTM α-synuclein

Fig. S2. Downstream consequences of NOX2-mediated α-synuclein modifications

Fig. S3. The NOX2 inhibitor, CPP11G, is blood brain barrier permeable, does not interfere with rotenone metabolism and inhibits NOX2 24 hours after an oral dose.

Fig. S4. NOX2 inhibition prevents rotenone-induced loss of body mass in rats.

Fig. S5. NOX2 in PD pathogenesis.

## Materials and methods

### Reagents

All reagents were purchased from Sigma-Aldrich, unless otherwise specified. Selective NOX2 kinase inhibitors Nox2ds-tat and CPP11G were provided by Dr. Patrick J. Pagano.

Antibodies were sourced and used as follows:

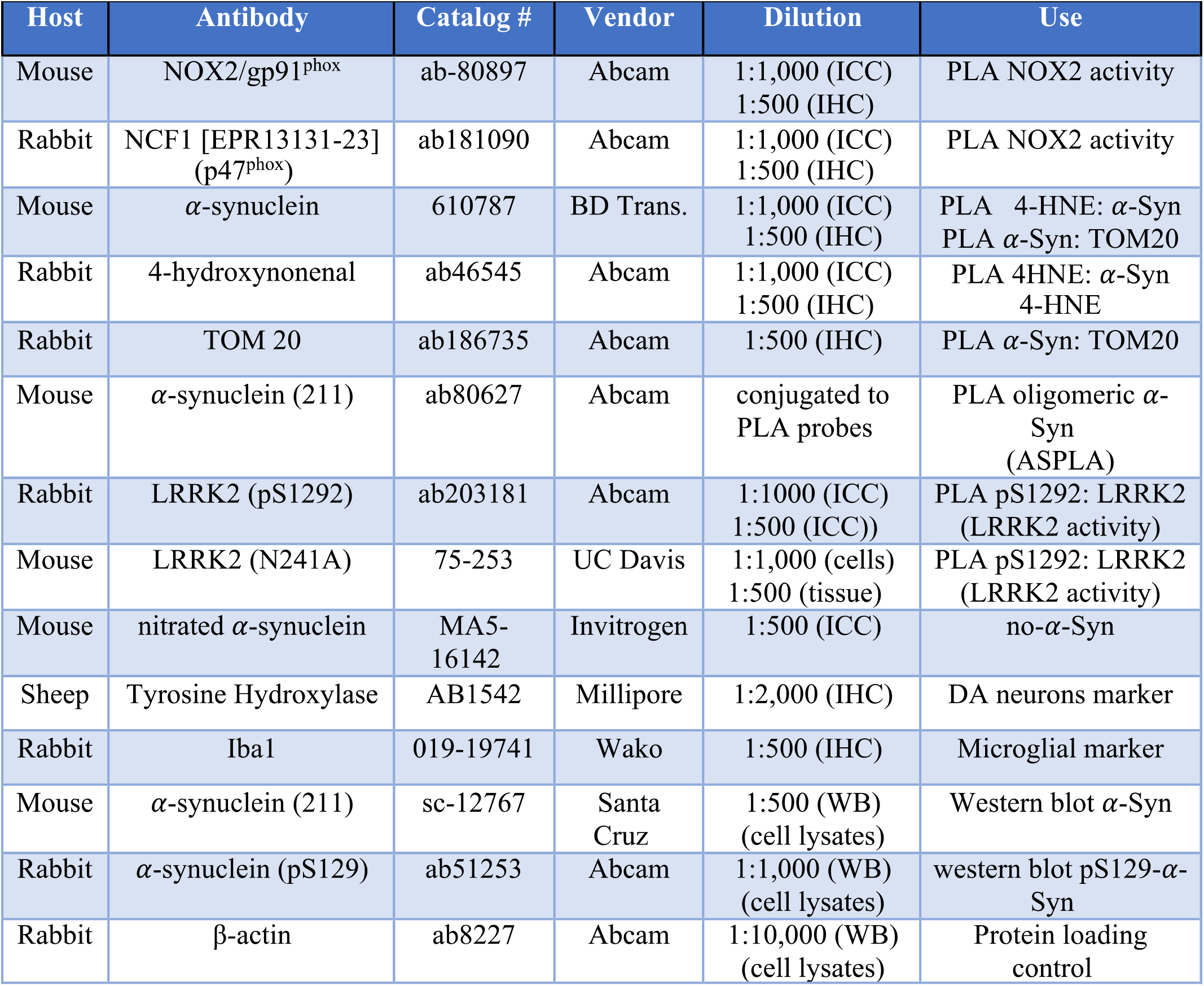

### Fluorescence measurements

Quantitative fluorescence measurements were made with an Olympus upright 3-laser scanning confocal microscope, taking care to ensure that images contained no saturated pixels. For quantitative comparisons, all imaging parameters (e.g., laser power, exposure, pinhole) were held constant across specimens. Depending on the specific experiment, readouts included fluorescence or number of objects (punctae) intensity in predefined regions of interest, such as tyrosine hydroxylase-positive dopaminergic neurons or Iba1-positive microglia.

### Animals

All experiments utilizing animals were approved by the Institutional Animal Care and Use Committee of the University of Pittsburgh. Rats were treated with rotenone as described (49, 50). For details of AAV2-mediated gene transfer, see (6, 7, 33, 50).

### AAV2-mediated gene transfer

Details per Zharikov et al (50). Rats were euthanized 6 weeks after injection.

### Human tissue

Paraffin-embedded midbrain sections were obtained from the University of Pittsburgh Brain Bank. All banked specimens have undergone standardized premortem neurological and postmortem neuropathological assessment. Diagnoses were confirmed and staging performed by the study neuropathologist (JKK) by examination of H&E, alpha-synuclein, tau, silver and ubiquitin stains of key sections needed for Braak staging (51). The study design was reviewed and approved by the University of Pittsburgh Committee for Oversight of Research Involving the Dead. Midbrain sections from 6 PD/PDD patients and 7 control subjects, matched for age and postmortem intervals, were used for analysis.

To eliminate endogenous fluorescence, human tissue was pre-treated with an autofluorescence eliminating reagent according to the manufacturer’s instructions (Chemicon, Temecula, CA).

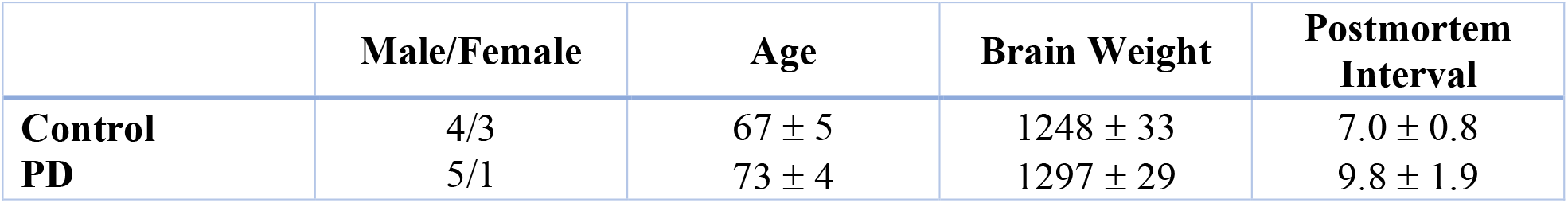

### Detection of CPP11G and rotenone concentrations in rat brains

1.5 pmol CPP11H internal standard was added to 50 μl of rat brains homogenates and 300 μl acetonitrile was added for extraction. After sonication and centrifugation, the supernatants were analyzed for CPP11G and rotenone by high performance liquid chromatography-electrospray ionization-tandem mass spectrometry. After chromatographic resolution, CPP11G and rotenone were detected using a 5000 triple quadrupole mass spectrometer (Applied Biosystems, San Jose, CA) equipped with an electrospray ionization source (ESI) in positive mode.

The following parameters were used for the analysis of CPP11G: declustering potential (DP) 80 V, entrance potential (EP) 10 V, collision cell exit potential (CXP) 15 V, and a desolvation temperature of 700 °C. Multiple reaction monitoring (MRM) transitions were used with the best collision energy (CE): MRM 390.1/334.1, 390.1/278 and 390.1/207.9 with CE 20 eV, 35 eV, and 45 eV respectively. For rotenone analysis: DP 80 V, EP 5 V, CE 32 eV, CXP 13 V, MRM 395.3/367.2, 395.3/213.2 and 395.3/192.2. Quantitation of CPP11G in rat brain was performed using CPP11G calibration curves in presence of CPP11H as internal standard (MRM 416.1/97). Levels of CPP11G in rat brain were reported as pmol/mg protein. Quantitation of rotenone in rat brain was performed using an external calibration curve and levels of rotenone were reported as pmol/mg protein.

**Supplementary Fig. 1.**
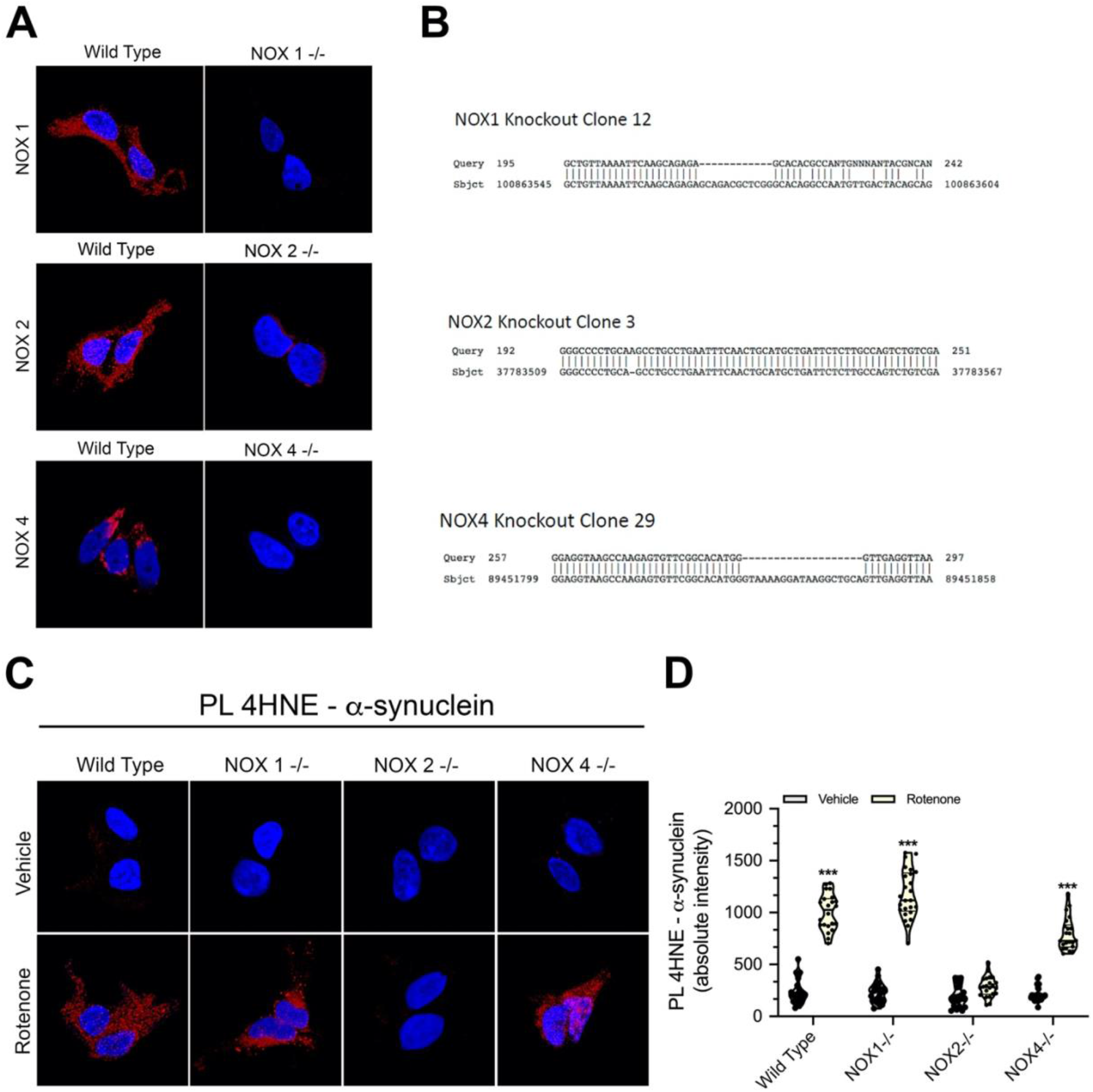
NOX2 is the key NADPH oxidase isoform in the induction of PTM α-synuclein formation. (**A**) Immunofluorescent staining for Nox1, Nox2 and Nox4 using isoform specific antibodies in wildtype cells (left column) and CRISPR/Cas9-edited Nox1^-/-^, Nox2^-/-^ and Nox4^-/-^ cells (right column) demonstrates absence of protein expression in the NOX isoform-specific null cells. (**B**) Sequence analysis shows deletions in NOX1and NOX4 and insertion in NOX2 genes supporting the knockout of the proteins. (**C**) PL assay for 4-HNE-modified α-synuclein (red). Only in NOX2^-/-^ cells was rotenone ineffective in inducing 4-HNE-α-syn adducts, indicating an exclusive role of NOX2 in the formation of this oxidative stress-related form of α-synuclein. (**D**) Quantification of 4-HNE-α-syn PL signal. Symbols represent the average of single cells fluorescence intensity analyzed in one 100X microscopic field. Statistical comparison by one-way ANOVA with post hoc Bartlett’s correction on three independent experiments. *** denotes p <0.0001 significance compared vehicle.

**Supplementary Fig. 2.**
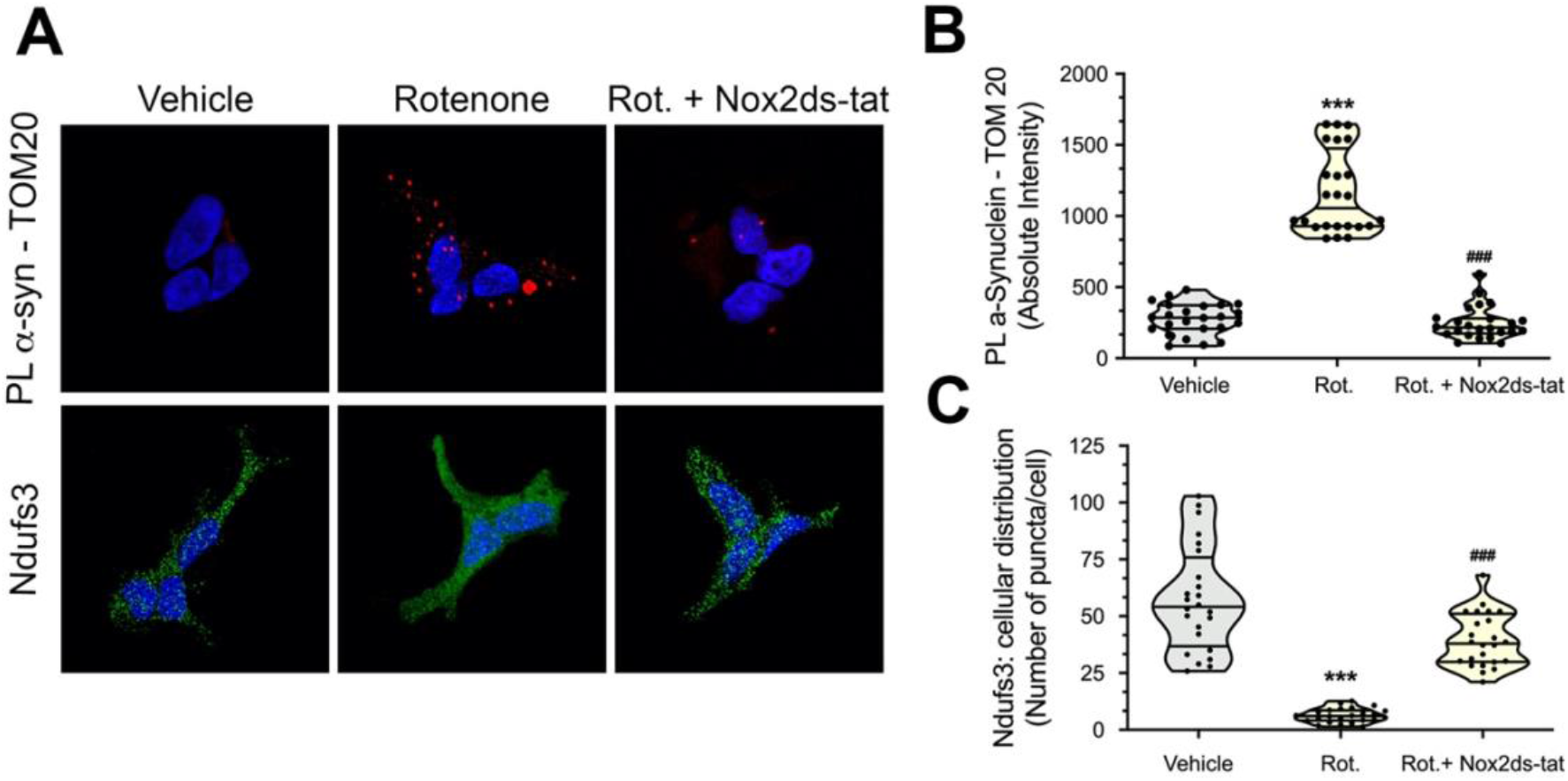
Downstream consequences of NOX2-mediated α-synuclein modifications. (**A**) *Top row* (red): PL α-syn-TOM20 interaction; and *bottom row* (green) Ndufs3 cellular distribution. Rotenone treatment for 24 hours enhanced the α-syn-TOM20 interaction (PL signal) in wild-type cells and re-distributed Ndufs3 from mitochondrial *punctae* to a diffuse cytosolic localization, consistent with impairment of mitochondrial protein import. Nox2ds-tat co-treatment prevented the α-syn-TOM20 interaction and Ndufs3 redistribution, suggesting a crucial role of NOX2 in α-synuclein-mediated mitochondrial dysfunction. (**B**) Quantification of α-syn-TOM20 PL signal. Symbols represent the average of single cells fluorescence intensity analyzed in one 100X microscopic field. Statistical comparison by one-way ANOVA with post hoc Bonferroni correction on three independent experiments. (**C**) Ndufs3 distribution analysis performed as number of puncta per cell. Symbols represent the average of single cells puncta counts analyzed in one 100X microscopic field. Statistical comparison by one-way ANOVA with post hoc Bartlett’s correction on three independent experiments. *** denotes p <0.0001 significance compared vehicle; ### denotes p <0.0001 significance compared rotenone.

**Supplementary Fig. 3.**
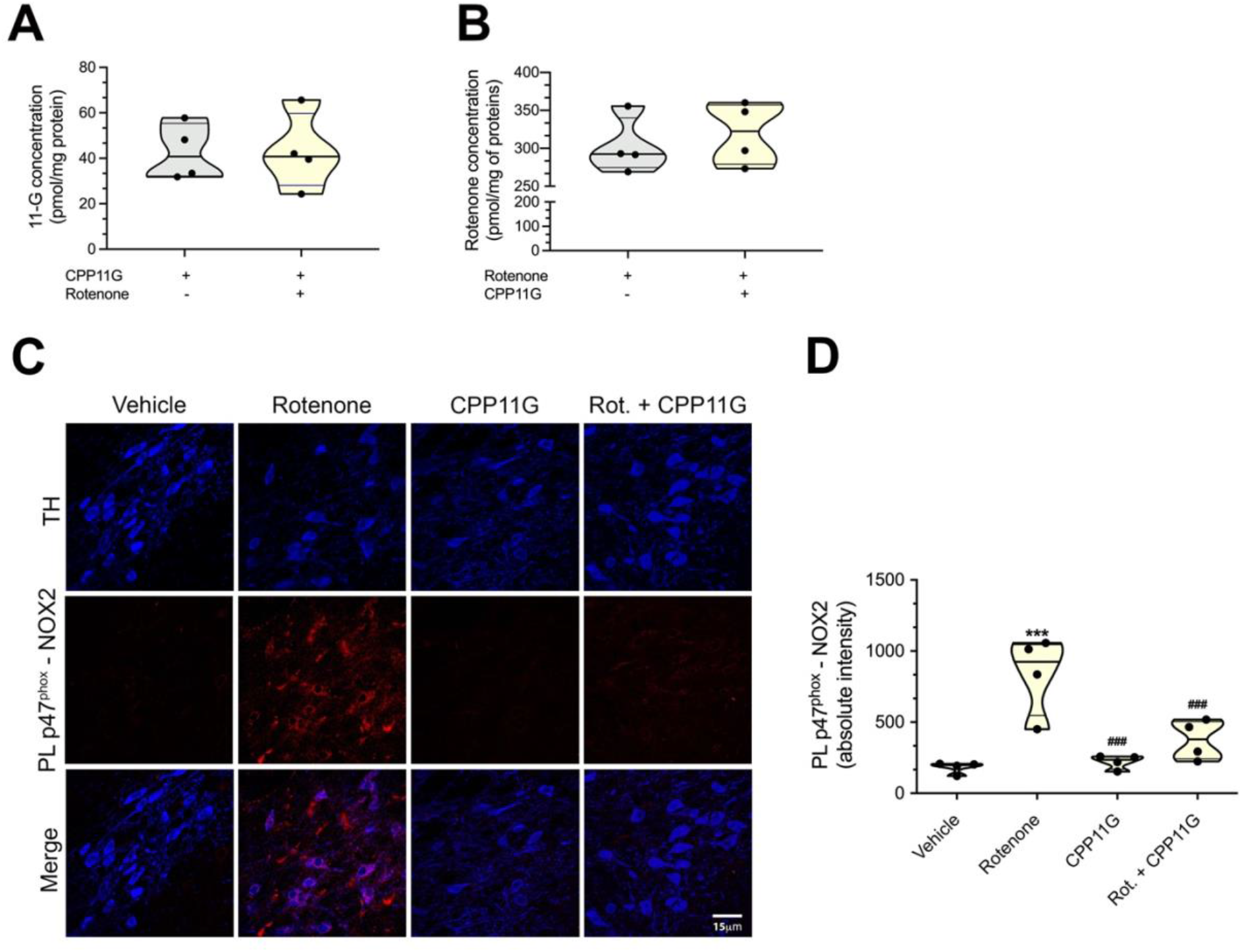
The NOX2 inhibitor, CPP11G, is blood brain barrier permeable, does not interfere with rotenone metabolism and inhibits NOX2 24 hours after an oral dose. **A**) Quantification of mass spectrometry analysis for CPP11G in brain lysates 24 hours after CPP11G administration (15 mg/Kg; gavage). Co-administration of CPP11G and rotenone (2.8 mg/Kg; i.p.) did not affect the residual brain concentration of CPP11G. (**B**) Mass spectrometry analysis of rotenone revealed no difference in brain concentration when co-administered with CPP11G. Each symbol represents individual animal’s brain lysate. Statistical comparison by paired two-tailed t test. (**C**) PL-based *in situ* detection of NOX2 activity status (second row, red) 24 hours after a single rotenone dose. Nigrostriatal neurons showed increased NOX2 activity detected as p47^*phox*^-NOX2 PL signal in dopamine neurons of substantia nigra (TH, blue). This was prevented by co-administration of CPP11G with rotenone, indicating that once daily administration of CPP11G (15 mg/Kg; gavage) inhibits NOX2 for at least 24 hours. (**D**) Quantification of PL signal for NOX2 activity. Each symbol represents individual animal. Statistical comparison by one-way ANOVA with post hoc Bartlett’s correction. *** denotes p <0.0001 significance compared vehicle; ### denotes p <0.0001 significance compared rotenone.

**Supplementary Fig. 4.**
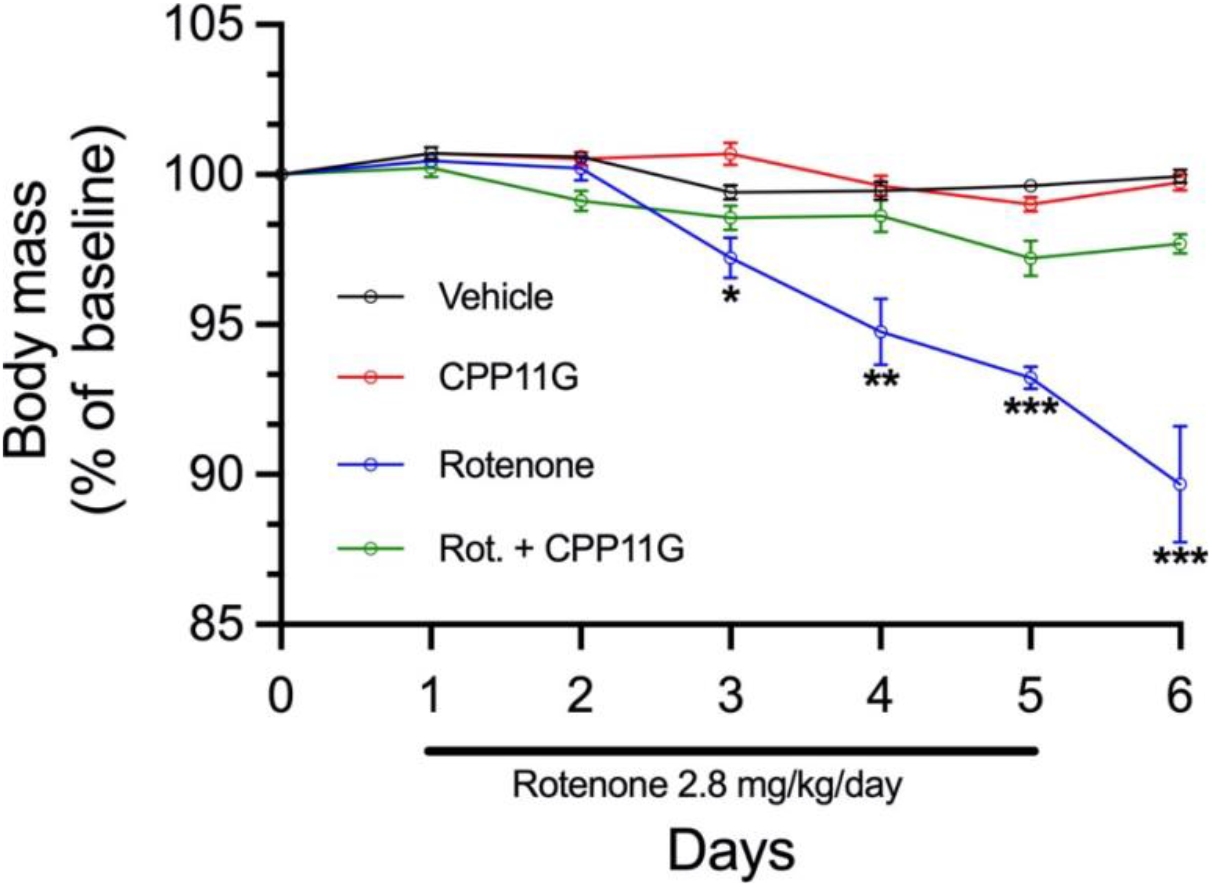
NOX2 inhibition prevents rotenone-induced loss of body mass in rats. Body mass was recorded daily in rats treated for 5 days with: vehicle, CPP11G alone, rotenone alone, or rotenone + CPP11G. Rotenone treatment caused about a 10% loss of body mass over 6 days, which was prevented by co-administration of CPP11G.

**Supplementary Fig. 5.**
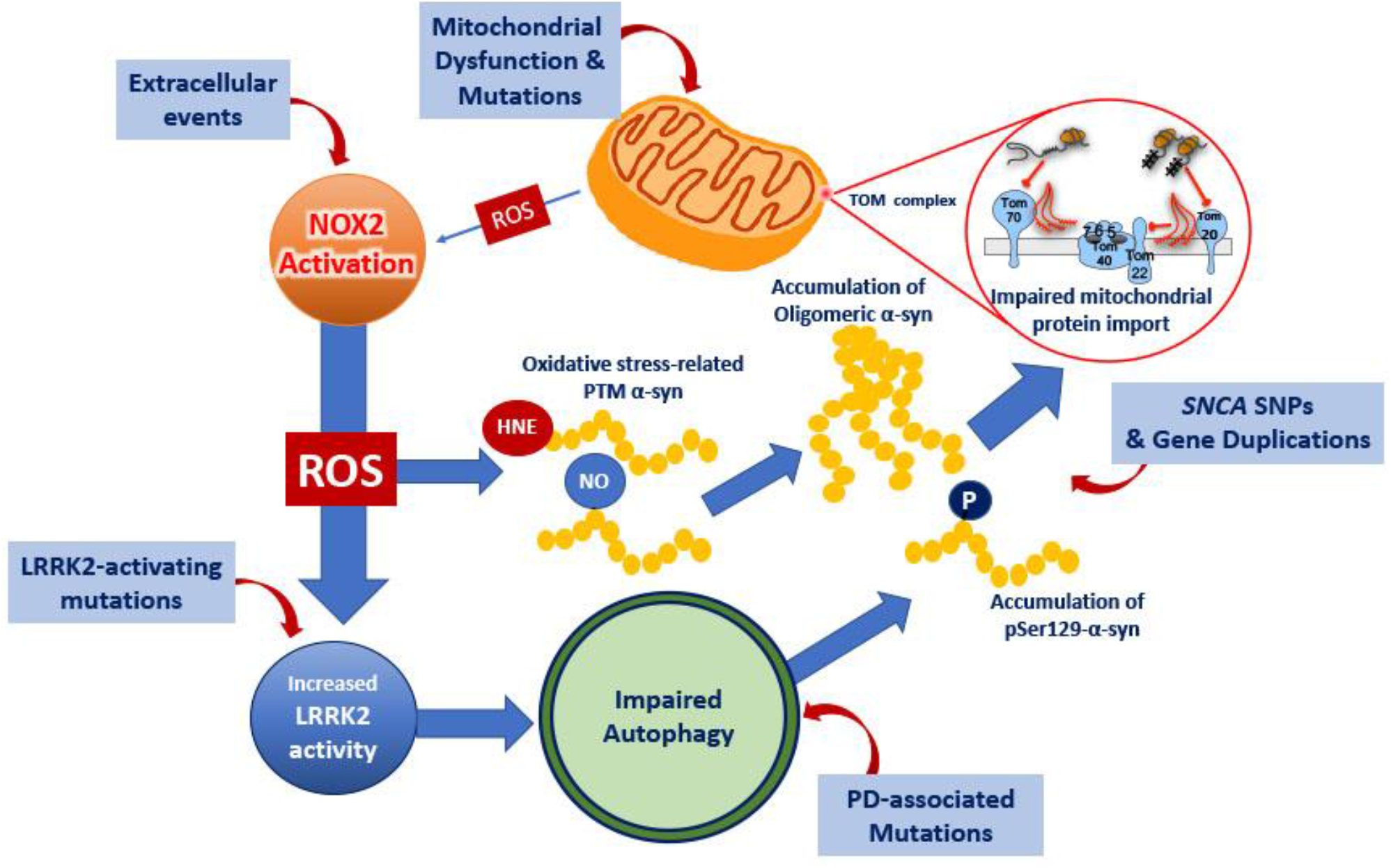
NOX2 in PD pathogenesis. The present study reveals that neuronal NOX2 can be activated by mitochondrially-derived ROS, in a process called ROS-induced ROS production, which serves to greatly amplify ROS production. The resultant NOX2 activity is necessary and sufficient to both activate wildtype LRRK2 and to cause accumulation and oxidative posttranslational modifications of α-synuclein (4-HNE-α-syn and NO-α-syn). The NOX2-activated LRRK2 activity leads to endolysosomal and autophagic defects and decreased degradation of toxic species of α-synuclein, such as pSer129-α-synuclein. In this way, NOX2 activity both drives the formation and inhibits the disposal of toxic forms of α-synuclein. In turn, as shown in previous work, these forms of α-synuclein bind to mitochondrial TOM20, blocking protein import and leading to mitochondria that produce more ROS. Thus, there appears to be a feedforward cycle, in which neuronal NOX2 plays an essential amplification role. Importantly, in this scheme, the cycle can be initiated at any of several nodes, including (1) environmental mitochondrial toxicants, such as rotenone, (2) mitochondrial mutations, such as parkin or Pink-1, (3) pathogenic LRRK2 mutations, (4) PD-associated mutations that affect endolysosomal function, or (5) *SNCA* polymorphisms or copy number variants that lead to increased levels of α-synuclein protein. In this context, it will be of great interest to determine whether inhibition of NOX2 can break this pathogenic cycle regardless of how it is initiated.

